# Xenogeneic Fibroblasts Inhibit The Growth Of Breast And Ovarian Cancer Cell Lines In Co-Culture

**DOI:** 10.1101/2021.07.19.452126

**Authors:** L. Usha, O. Klapko, S. Edassery

**Author notes:** Corresponding author: Lydia Usha, M.D., Division of Hematology, Oncology, and Stem Cell Transplant, Department of Medicine, 1725 W. Harrison St, Suite 809, Chicago, IL 60612. Northwestern University I Feinberg School of Medicine, Department of Neurology, Chicago, IL 60611.

## Abstract

Cell-based therapies cure some hematologic malignancies, although little information exists on solid cancer cell responses. The study objective was to test the hypothesis that xenogeneic fibroblasts can inhibit the growth of human cancer cell lines *in vitro*. Seven human cell lines (pancreatic cancer HPAF II; brain cancer U-87 MG; fibrosarcoma; ovarian cancer OVCAR3 and SKOV3; and breast cancer MCF7 and MDA-MB231) were co-cultured with two xenogeneic fibroblast cell lines (CV-1;monkey, *Cercopithecus aethiops* and DF-1; chicken, *Gallus gallus*) in a transwell culture system. Cancer cell proliferation was assessed colorimetrically. Different concentrations of breast and ovarian cancer cells were tested. Gene expression induced by DF-1 xenogeneic fibroblasts was assessed by RNAseq of MCF7 breast cancer cells. The proliferation of the majority of the cancer cell lines was altered by co-culture with xenogeneic fibroblasts. Cell proliferation was increased (4-17%) by CV-1; DF-1 increased brain cancer cell proliferation (16%), decreased breast and ovarian cancer cell growth (15 and 26% respectively) but did not affect fibrosarcoma and pancreatic cancer cells. When the initial cancer cell concentrations were lowered 4-fold, growth inhibition of breast and ovarian cancer increased more than 2-fold. DF-1 fibroblasts induced significant differential expression of 484 genes in MCF7 breast cancer cells; 285 genes were down-regulated and 199 genes were upregulated compared to control. Genes involved in the immune response were the major downregulated entities. RNAseq results were validated by qRT-PCR of 12 genes. The results show that xenogeneic fibroblasts can alter the growth and gene expression of cancer cells *in vitro*. This suggests a potentially novel investigational approach to the control of cancer cell growth.

## INTRODUCTION

Metastatic cancer remains a lethal disease despite numerous advances in cancer treatment. Although existing cancer treatments such as surgery, radiotherapy, and chemotherapy can achieve a cure in some early-stage cancers, they have only a palliative effect in advanced cancers. In addition, they often have complications and toxicities. For some malignancies with a single gene defect, treatment progress and sometimes a cure is achieved by “targeted” drugs such as the PARP inhibitor olaparib for ovarian cancer in women with germline BRCA mutations [1, 2] and imatinib which targets BCR-ABL tyrosine kinase in Philadelphia chromosome-positive chronic myelogenous leukemia [3]. However, in the majority of cases, the common epithelial malignancies (e.g., colon, breast, lung, liver, pancreas and ovary) have multiple genetic alterations that often involve deletion and amplification of large parts of the genome, as well as complex and interacting cellular and molecular networks. Consequently, the success of single-target therapies or even their combinations is often limited.

A potential alternative to targeted therapies is cytotherapy. Cytotherapy has been successfully used to treat cancer. Although commonly used in experimental conditions cells include mesenchymal stem cells (MSC) and fibroblasts, a variety of other cells were investigated [4, 5]. For example, CAR-T (chimeric antigen receptor) cell therapy was recently developed and FDA-approved for some forms of leukemia [6].

Xenogeneic cytotherapies have shown some promise. Xenogeneic cells have the potential to stimulate the immune system to overcome tumor immunosuppression [7-9]. An influx of immune cells and tumor regression occurred in patients with solid metastatic tumors in a Phase 1 clinical trial in which xenogeneic African green monkey fibroblasts, engineered to produce human IL-2, were administered intratumorally [10]. Likewise, targeted cytotherapy for pancreatic cancer using naïve, non-engineered rat umbilical cord matrix derived stem cells to control the growth of pancreatic cancer, strongly attenuated the growth of pancreatic carcinoma cells *in vitro* and *in vivo* in a peritoneal mouse model [11]. Xenogeneic immunization with tyrosine hydroxylase-derived DNA vaccines was effective against neuroblastoma in mice [12]. In a human clinical trial performed in Russia, irradiated xenogeneic murine cell vaccines were effective in breaking the immune tolerance to human tumor-associated antigens in human colorectal cancer [13].

One appeal of cell therapy is that cells may express and secrete thousands of biologically active molecules which could interfere with malignant growth. Also, specific cells, such as stem cells or fibroblasts, can migrate to the corresponding tissue/organ (“homing”) or at least in the case of mesenchymal stem cells, to the site of injury, inflammation or cancer [14, 15]. Once “homed” to the tumor, xenogeneic fibroblasts can potentially elicit a robust immune response to the tumor. Cancer-associated fibroblasts play a fundamental role in modulating the behavior of cancer cells and cancer progression [15, 16]. There is evidence for both pro- and anti-tumor actions of cancer-associated fibroblasts [17]. Interestingly, normal human and murine fibroblasts inhibit the proliferation and motility of prostate tumor cell lines when co-cultured *in vitro* [18]. This effect involved both cell contact with fibroblast monolayers and fibroblast secreted biomolecules.

The objective of the study was to assess the effect of xenogeneic fibroblasts on cancer cells in order to test the hypothesis that xenogeneic fibroblast cell lines can inhibit the growth of human cancer cell lines in vitro. The approach was to use a simplified system of exposing cancer cells to xenogeneic fibroblasts *in vitro* in which cells are separated in a transwell system. Cancer cell growth and molecular expression changes in cancer cells were examined.

## METHODS

### Cell Culture

All cell lines were obtained from the American Type Culture Collection (ATCC; Manassas, VA). The two fibroblast cell lines included UMNSAH/DF-1 (henceforward designated DF-1) derived from normal *Gallus gallus (chicken)* embryo fibroblasts (ATCC CRL12203) and CV-1 derived from normal *Cercopithecus aethiops* (African green monkey) kidney fibroblasts (ATCC CCL70). The seven human cancer cell lines included those from pancreatic cancer HPAF II (ATCC CRL1997); brain cancer U-87 MG (ATCC HTB14); fibrosarcoma (ATCC CCL121); ovarian cancer OVACAR3 (ATCC HTB161) and SKOV3 (ATCC HTB77); and breast cancer MCF7 (ATCC HTB22) and MDA-MB231 (ATCC HTB26). After experiments at a fixed starting cancer cell concentration, the two breast cancer cell lines and two ovarian cancer cell lines were selected to explore the effects of initial cancer cell concentrations on growth effects in co-culture with the chicken embryo-derived fibroblast cell line. The breast cancer cell lines were estrogen and progesterone receptor-positive (MCF7) or estrogen and progesterone-negative (MDA-MB231); ovarian cancer cell lines included a p53 mutant cell line (OVCAR3) and a p53 wild-type cell line (SKOV3).

Cells were cultured in Eagles Essential minimum media (ATCC) containing 10% fetal bovine serum (FBS) (Sigma; St. Louis, MO) in a humidified incubator (5% CO_2_) at 37°C. Cancer cells were co-cultured with fibroblast cell lines using a transwell system (Greiner Bio-one Thincert^™^; Monroe, NC) with fibroblasts layered in the upper well (Figure 1). Cancer cells were seeded at 4⨯10^4^ cells/well/800 µl of media and fibroblast cells in the insert at 2×10^4^ cells/300 µl media. As a control, the matching cancer cells were used in both the well and insert with the cancer cells in the insert at 2⨯10^4^ cells/300 µl media.

**Figure 1.**
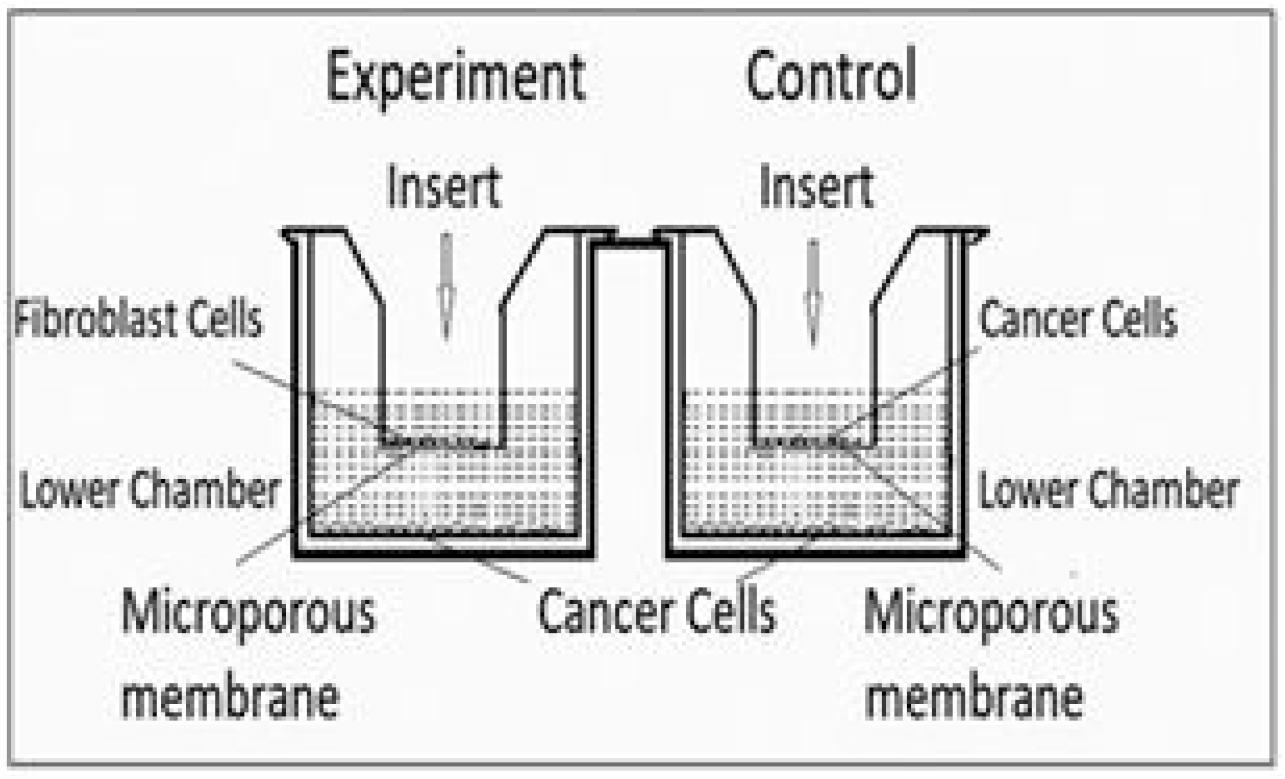
Diagram of transwell used for co-culture of fibroblasts and cancer cell lines Legend: In the experimental wells, cancer cells were seeded in the lower chamber and fibroblasts were seeded in the upper well. In the control wells the same cancer cells were seeded in the top and bottom wells without added fibroblast cell lines. Upper wells contained 2×10^4^ cells/300 µl media.

Cultures were monitored for mycoplasma contamination. Also, RNA sequencing data were analyzed for mycoplasma sequences. The data were uploaded into the Galaxy server, four million reads from each file were selected using the “split file tool,” and the split FASTQ sequencing file aligned using “Bowe2” against three mycoplasma fasta sequences (*M. fermentans* M64, *M. hominis* ATCC 23114 and *M. hyorhinis* MCLD) commonly found in cell cultures [19]. No alignment with the mycoplasma sequences was found, supporting a lack of mycoplasma contamination.

### Cell Growth

After five days, the growth of the cancer cells was measured using the MTT (3-(4,5-dimethylthiazol-2-yl)-2,5-diphenyltetrazolium bromide) cell proliferation assay. The reduction of yellow tetrazolium dye (MTT) by living cells produces purple formazan crystals. In brief, the inserts were removed, and 80µl of 5 mg/ml MTT reagent (MP Biomedicals; Santa Ana, CA) was added to each well and incubated for 4 hours. The media was removed, and the intracellular formazan solubilized by adding 500 µl of DMSO (Sigma; St. Louis, MO) and incubating for 30 minutes at room temperature. Optical density was read at 560 nm with a reference wavelength of 650 nm using an Epoch spectrophotometer (BioTek Instruments; Winooski, VT). The percent change in growth compared to the control incubations was calculated. The experiment was repeated three times for cell growth assays and RNA extraction. The data were analyzed using the Student’s t-test with p< 0.05 considered significant.

### RNA Extraction

The effect of the DF-1 fibroblast cell line on gene expression changes in human MCF7 breast cancer cells was determined. The media was removed after co-culture for five days, and the culture wells were washed with cold phosphate-buffered saline (PBS). RNA was extracted with 100 µl of Trizol per well, triplicate wells were pooled, and the Trizol extractions continued according to the manufacturer’s protocol. Precipitated RNA was solubilized in 40 µl of molecular biology grade water (RPI Research Products; Mount Prospect, IL) and stored at -80°C. RNA quantity and quality was measured in a NanoDrop Spectrophotometer. Novogene checked RNA integrity with an Agilent 2100 analyzer, and RNA degradation was assessed by gel electrophoresis before subjecting the sample to Next-Generation sequencing.

### Next-Generation Sequencing

The extracted RNA was sent to Novogen Inc. (Davis, CA) for sequencing. cDNA libraries were made from the RNA as per the in-house Novogene protocol. Paired-end (PE-150bp) sequencing was performed using the Illumina sequencing platform; each RNA sample was sequenced to obtain at least 40 million reads per sample. The RNA sequence data was “cleaned” using the NOVOGENE default protocol to remove the adapter sequences, reads containing more than 10% N (sequence not determined) and reads with a low-quality Phred score (Q Score <=5) (Supplementary Table 1).

The cleaned data was uploaded to Galaxy (https://usegalaxy.org/) for analysis. Sequence count per gene was determined using the “Salmon” method in Galaxy. The Salmon program quantifies the expression of transcripts from RNAseq data; it indexes, quantifies, and provides the count per each transcript aligned to the reference genome [19]. The count files from each sample are merged into one list and uploaded to the Degust webserver for differential gene expression (DE) analysis between groups. The data was further filtered to remove “no count” transcripts by using criteria of at least one count per transcript in all samples. EdgeR implemented in the Degust (http://degust.erc.monash.edu/) was used to identify differentially expressed genes (FDR = 0.01 and absolute log fold change = +/- 0.5), using the quasi-likelihood functionality of edgeR; DE genes lists were analyzed further for functional enrichment using the “Reactome pathway browser “at https://reactome.org/PathwayBrowser/.

### qRT-PCR

Examples of differentially expressed genes were selected for confirmation and validation: interleukin 1 receptor, type I, cysteine-rich secretory protein 3, KIT ligand, selectin L, G protein-coupled estrogen receptor 1, protein kinase C, delta, interleukin 1 receptor antagonist, chemokine (C-X-C motif) ligand 12, CD36 molecule (thrombospondin receptor), B-cell CLL/lymphoma 2, Janus kinase 2, and interleukin 18. The qRT PCR primers were designed using NCBI primer blast. One microgram of total RNA was treated with DNASE (Thermo Fischer; Waltham, MA) to remove any genomic DNA contamination according to the manufacturer’s standard protocol. Five hundred ng of DNASE treated RNA was used for the first-strand synthesis using an ABI high-capacity cDNA reverse transcription kit (Applied Biosystems; Foster City, CA). qRT-PCR was carried out using Fast Sybr green (ABI). Ct values were exported, and the ddCt value was used to calculate fold change in differential expression.

## RESULTS

### Cell growth

The proliferation of the majority of the cancer cell lines was altered by transwell co-culture with xenogeneic fibroblast cell lines.

Specifically, CV-1 fibroblasts increased the proliferation of brain cancer cells by 12%, OVCAR3 ovarian cancer cells by 13.5% and fibrosarcoma cells by 11% compared to control. These changes were significant (p<0.05). The growth of the pancreatic cancer cell line increased 20% in two out of three experiments, while growth of the MCF7 breast cancer cell line did not increase significantly (Figure 2A).

**Figure 2.**
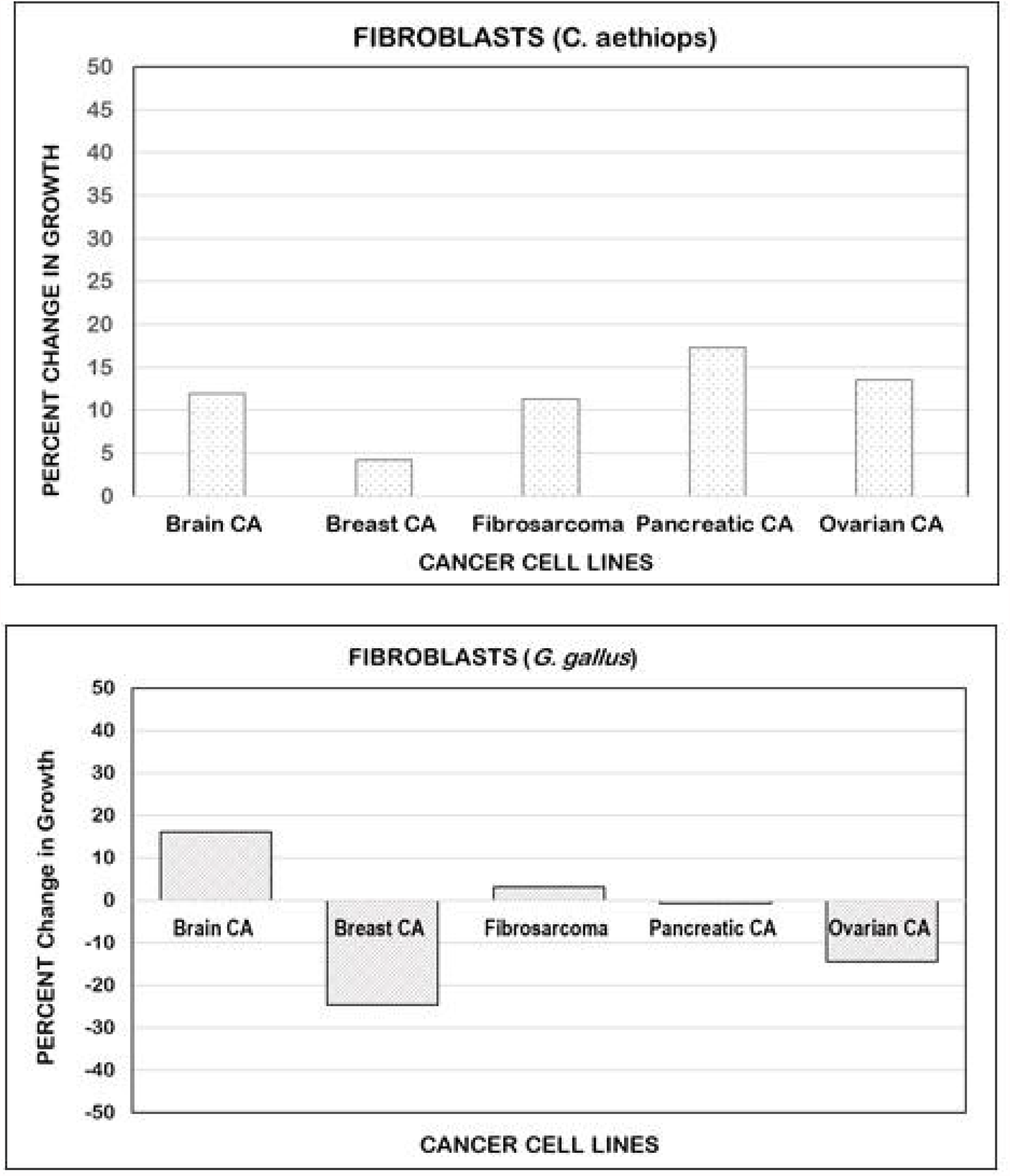
Xenogeneic fibroblast cell lines alter cancer cell growth in co-culture Legend: The proliferation of a majority of the cancer cell lines was increased significantly (p<0.05) in the presence of CV-1 fibroblasts (monkey; *C. aethiops)* (upper panel). The percent change in growth is shown for brain cancer (HTB14 cells), ovarian cancer (HTB161 line of OVACR3 cells), fibrosarcoma (CCL121 cells) and pancreatic cancer (HPAF II and CRL1997 cells) compared to the control. The small increase in the breast cancer cell line MCF7 (HTB22) growth was not significant. The DF-1 (chicken; *G*.*gallus)* embryo-derived fibroblast cell line had mixed effects on the cancer cell lines (lower panel); the growth of brain cancer increased but the growth of breast cancer MCF7 (HTB22) and OVCAR3 was reduced compared to the control (p<0.05). The pancreatic cancer cell line (HPAF II, CRL1997) and the fibrosarcoma cell line (CCL121) showed no significant change in response to DF-1 cells. All experimental wells were seeded at 4⨯10^4^ cancer cells/well/800 µl of media along with 2⨯10^4^ cells/300 µl media fibroblast cells in the transwell insert. Controls consisted of adding matched cancer cells to the transwell.

DF-1 fibroblasts had mixed effects on cancer cell line growth. Brain cancer cell proliferation increased 16%. The growth of MCF7 breast cancer cells was reduced 25% and OVCAR3 ovarian cancer cell growth was reduced by 14% compared to control (p<0.05). The pancreatic cancer and the fibrosarcoma cell lines showed no significant growth change in co-culture with either fibroblast cell line (Figure 2B).

To assess the effect of different initial cancer cell concentration (10,000, 20,000 and 40,000) on cell growth in the presence of xenogeneic fibroblasts, proliferation was examined for the breast and ovarian cancer cell lines in co-culture with DF-1 fibroblasts. Inhibition was higher overall at lower starting cell concentrations (Figure 3) except for OVCAR3 where inhibition was higher at the highest cell concentration. Thus the growth effects differed between the two ovarian cancer cell lines. The pattern was similar for either breast cancer cell lines regardless of estrogen and progesterone receptor absence (e.g., MDA-MB231 cells) or presence (e.g., MCF7 cells).

**Figure 3.**
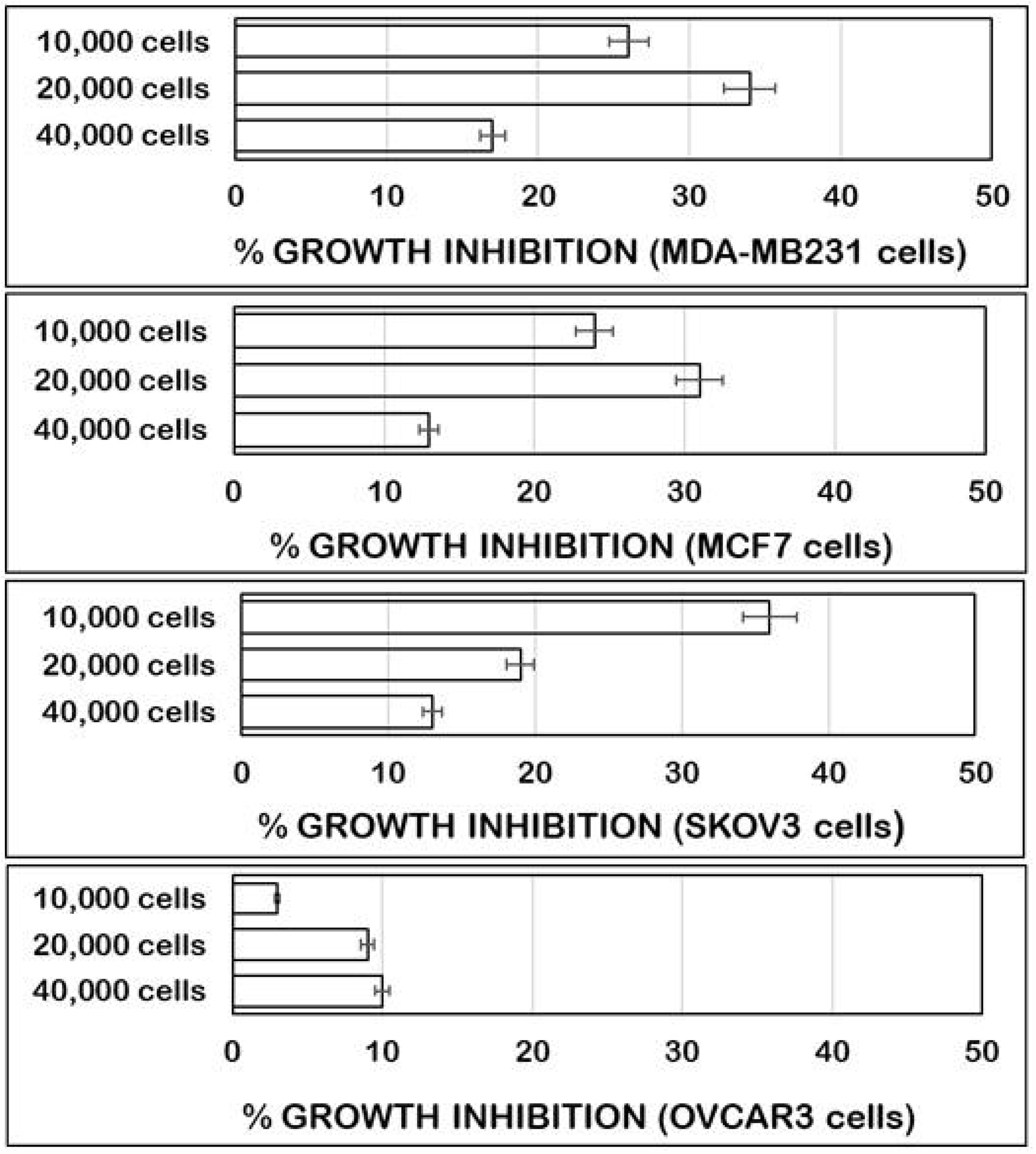
Cancer cell concentration and inhibition of their growth by chicken embryo-derived fibroblast Legend: Inhibition of cancer cell line growth by DF-1, (chicken; *G*.*gallus)* embryo-derived fibroblast cell line, is greater at lower initial cancer cell numbers.

### Gene Expression

Gene expression was examined in the MCF breast cancer cell line in response to DF-1 fibroblasts. Overall, 484 genes were significantly differentially expressed; of these, 285 genes were down-regulated and 199 genes were up-regulated compared to control. The results of Reactome analysis show the major functional pathways that are differentially up-regulated (Table 1A) and down regulated (Table 1B). Immune pathway genes (n = 101 entities found) were the major over-represented group among the downregulated genes. Among the upregulated genes those involved in developmental pathways were the most overrepresented genes (n=27 entities found). An MA plot of the distribution of significantly up and down-regulated genes is shown in Figure 4.

**Table 1.** Selected genes in MCF7 breast cancer cell line with altered expression after exposure to DF-1 fibroblasts Selected genes from Reactome analysis with altered expression by RNAseq of the MCF7 breast cancer cell line in response to DF-1 xenogeneic fibroblasts are shown. There were more (B) downregulated (n=285) than (A) upregulated (n=199) entities. The top 10 with up- or downregulated expression are shown. Overall the p values were more significant for downregulated genes. The major changes in expression in the cancer cells involved the immune system.

| Pathway ID | Pathway name | Entities |  |  |  |
| --- | --- | --- | --- | --- | --- |
|  |  | # found | total | p Value | FDR |
| R-HSA-428540 | Activation of RAC1 | 3 | 15 | 0.001 | 0.227 |
| R-HSA-1266738 | Developmental Biology | 27 | 1176 | 0.005 | 0.292 |
| R-HSA-5617472 | Activation of hindbrain anterior HOX genes during early embryogenesis | 6 | 116 | 0.005 | 0.292 |
| R-HSA-5619507 | Activation of HOX genes during differentiation | 6 | 116 | 0.005 | 0.292 |
| R-HSA-8985586 | SLIT2:ROBO1 increases RHOA activity | 2 | 8 | 0.006 | 0.292 |
| R-HSA-8937144 | Aryl hydrocarbon receptor signaling | 2 | 8 | 0.006 | 0.292 |
| R-HSA-428543 | Inactivation of CDC42 and RAC1 | 2 | 12 | 0.012 | 0.490 |
| R-HSA-2559585 | Oncogene Induced Senescence | 3 | 42 | 0.020 | 0.508 |
| R-HSA-419037 | NCAM1 interactions | 3 | 44 | 0.023 | 0.508 |
| R-HSA-9018681 | Biosynthesis of protectins | 2 | 18 | 0.025 | 0.508 |

**B. Downregulated Genes**
| Pathway ID | Pathway name | Entities |  |  |  |
| --- | --- | --- | --- | --- | --- |
|  |  | # Found | Total # | P value | FDR |
| R-HSA-983170 | Antigen Presentation: Folding, assembly and peptide loading of class I MHC | 56 | 102 | 1.11E-16 | 6.99E-15 |
| R-HSA-1280215 | Cytokine Signaling in Immune system | 81 | 1051 | 1.11E-16 | 6.99E-15 |
| R-HSA-913531 | Interferon Signaling | 67 | 388 | 1.11E-16 | 6.99E-15 |
| R-HSA-198933 | Immunoregulatory interactions between a Lymphoid and a non-Lymphoid cell | 56 | 316 | 1.11E-16 | 6.99E-15 |
| R-HSA-1280218 | Adaptive Immune System | 62 | 999 | 3.20E-10 | 1.82E-08 |
| R-HSA-9018519 | Estrogen-dependent gene expression | 20 | 153 | 8.08E-09 | 4.28E-07 |
| R-HSA-8939211 | ESR-mediated signaling | 20 | 160 | 1.67E-08 | 8.04E-07 |
| R-HSA-9006931 | Signaling by Nuclear Receptors | 23 | 230 | 7.93E-08 | 3.57E-06 |
| R-HSA-168256 | Immune System | 101 | 2638 | 3.74E-05 | 0.001 |
| R-HSA-1474228 | Degradation of the extracellular matrix | 12 | 148 | 6.72E-04 | 0.021 |

**Figure 4.**
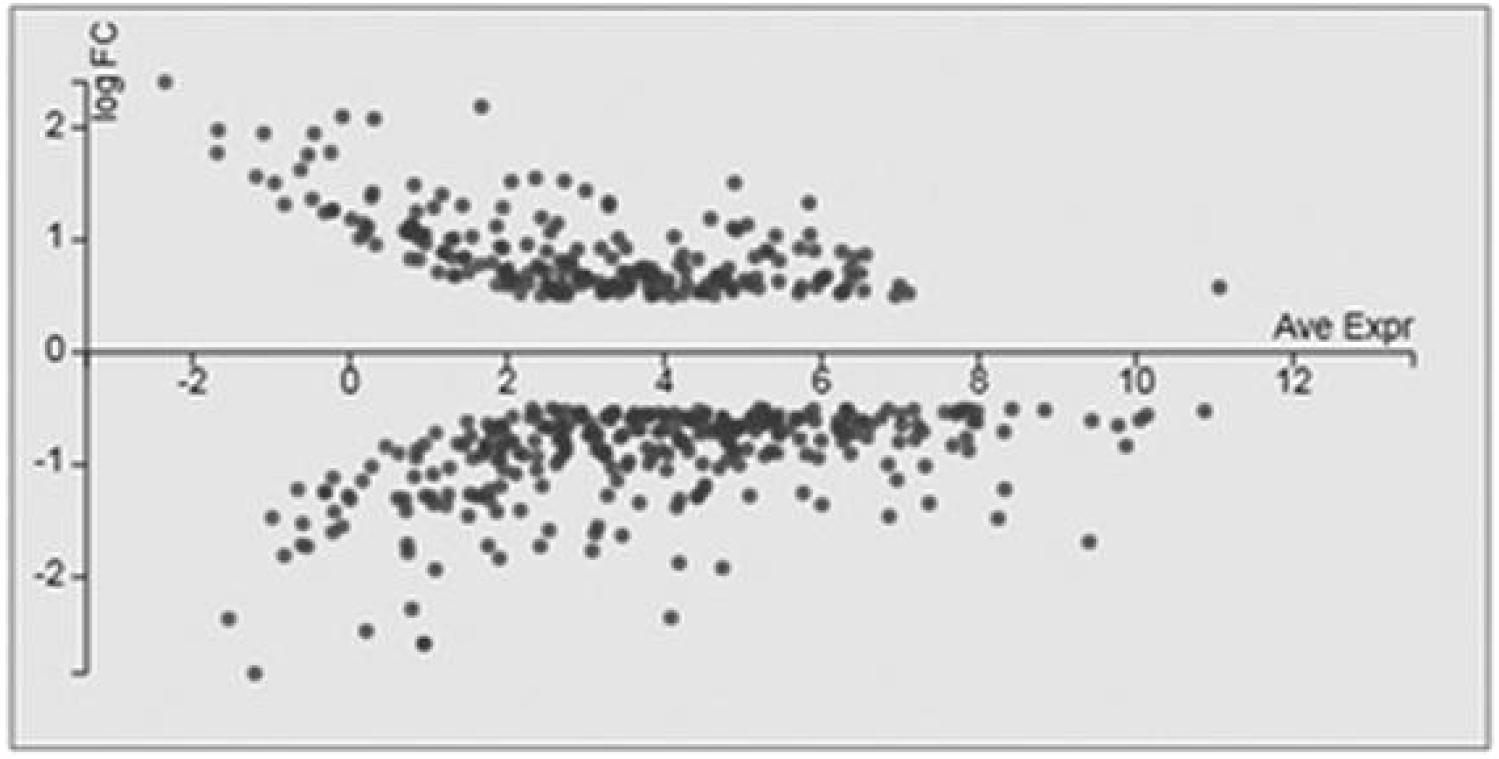
MA plot of significantly up- and downregulated genes in response to DF-1 fibroblasts Legend: MA plot shows the distribution of significantly (p<0.05) up- and downregulated genes in response to DF-1 xenogenic fibroblasts as a fold change (FC) in expression.

To validate the RNA sequencing data, 12 genes with a significant fold change in expression were chosen and quantified by qRT-PCR. The results of qRT-PCR supported those obtained from transcriptome analysis and demonstrated a similar up- or down-regulation of the genes (Table 2).

**Table 2.** The difference in expression of selected genes in MCF7 breast cancer cell line after exposure to DF-1 fibroblasts Changes in expression of selected genes seen in next generation sequencing (NGS) were confirmed by qPCR.

| Gene | NGS fold change | qRT PCR fold change |
| --- | --- | --- |
| interleukin 1 receptor, type I | 2.86 | 2.93 |
| cysteine-rich secretory protein 3 | 1.72 | 2.21 |
| KIT ligand | 1.64 | 2.20 |
| selectin L | 1.52 | 1.62 |
| G protein-coupled estrogen receptor 1 | 1.49 | 1.48 |
| protein kinase C, delta | 1.41 | 1.72 |
| interleukin 1 receptor antagonist | -3.13 | -1.75 |
| chemokine (C-X-C motif) ligand 12 | -2.27 | -1.75 |
| CD36 molecule (thrombospondin | -1.92 | -1.77 |
| B-cell CLL/lymphoma 2 | -1.89 | -1.74 |
| Janus kinase 2 | -1.79 | -1.60 |
| interleukin 18 | -1.79 | -1.60 |

## DISCUSSION

The proliferation of a majority of the cancer cell lines was altered by co-culture with xenogeneic fibroblasts. The direction of the effect varied with different cell combinations. The DF-1 chicken embryo-derived fibroblast cell line most effectively inhibited breast and ovarian cancer cell lines. Associated genomic changes in breast cancer cells showed that in addition to the expected changes in proliferation and differentiation pathways, downregulation of components of immune system pathways was the most dramatic change of the cancer cells.

These findings are significant since cancer-associated fibroblasts are an active component of the tumor microenvironment and they coordinate interactions between the cancer cells and stromal cells [15, 17]. Fibroblasts are known to remodel tumor stroma and can have pro- and anti-tumor effects [16, 21]. In this study, the difference between monkey and chicken fibroblasts effects on cell growth could be due to a number of factors. Aside from the obvious difference in species origin, the monkey fibroblasts are from adult kidney while chicken fibroblasts were from embryos that are pluripotent and could produce a different and greater range of active factors.

Interestingly, exposure to embryonic microenvironment(s) of chicken or zebrafish reprograms human melanoma cells and inhibits tumor development [22]. Fibroblasts are a dominant component of tumor stroma and play a key role in regulating the anti-tumor immune response [5]. Thus, the results of RNA-seq in this study are consistent with a major effect on immune system pathways.

Immunotherapy became an important new development in treatment of multiple malignancies and after more than 100 years of basic research and clinical trials, finally proved that manipulating the host immune system can lead to a clinically significant anti-tumor effect. However, the percentage of patients with advanced solid tumors who achieve a durable response or cure from immunotherapy remains small [23]. Immunotherapy employs monoclonal antibodies to biomarkers present on immune cells such as PD-L1 or CTLA-4 [23], and assumes all relevant targets are addressed. A potential advantage of cell therapy is that multiple factors are produced by therapeutic cells, which could address multiple targets.

Xenogeneic cell and organ transplantation has been used to replace damaged cells in Parkinson’s disease (DA neurons), diabetes (islets) and liver and heart failure [7]. Xenogeneic cell transplantation has been proposed as a therapeutic approach to re-activate anti-tumor immunity and restore impaired function [7]. Various cell types and preparations have been used as a cancer therapy [8]. Human, mouse and rat MSCs transfected with various vectors (e.g. viruses, transposon-based gene vectors) attenuated growth of different tumors targeted by each individual genetically engineered group of MSCs [24]. A composite xenogeneic polyantigenic vaccine prepared from murine melanoma B16 and carcinoma LLC cells increased survival and tumor immunity (e.g., increased T cell responses to human Caco-2 colon adenocarcinoma-associated antigens) in stage IV colorectal cancer patients [13]. Xenogeneic monkey fibroblasts (Vero cells) genetically engineered to produce human IL-2 were administered intra-tumorally; this treatment was associated with an anti-tumor effect [10]. A vaccine using xenogeneic whole endothelial cells effectively inhibited tumor growth, induced regression of established tumors and prolonged survival of tumor-bearing mice [9]. Fibroblasts inhibited cancer cell growth in co-culture; this inhibition varied depending on the source and the site of origin of the fibroblasts [18].

Caveats and considerations for this study are related to the heterogeneity of fibroblast functions. A uniform effect of fibroblasts was not expected since subsets of cancer associated fibroblasts have cancer-suppressing or cancer-promoting functions [15, 16]. Furthermore, fibroblasts have intrinsic cellular plasticity and exhibit heterogeneity in tumors [25]. Activated fibroblasts, which have similar features to mesenchymal stromal cells, may be capable of reprogramming into different lineages, including endothelial cells, adipocytes, and chondrocytes [17]. The function of the co-cultured fibroblasts may differ when paired with different cell lines. A future study would reveal the effect of fibroblasts in direct contact with cancer cells in co-culture.

Finally, this simplified system was used to gain knowledge of the effect of xenogeneic fibroblasts on cancer cells. However, it does not include the complex interactions that might occur *in vivo*; in other words, would the chicken fibroblast cell line drive the same response *in situ*? While there is interest in the development of fibroblast-targeted cancer therapies, the complexities of fibroblast-cancer cell interactions remain to be adequately understood before novel therapies targeting or exploiting fibroblasts can be implemented [5].

## CONCLUSION

The results of the studies showed that xenogeneic fibroblasts can inhibit proliferation of human ovarian and breast cancer cells *in vitro*. Furthermore, this growth inhibition was associated with differential expression of genes and genomic pathways involved in cell proliferation, differentiation, and immune function. Thus, the co-culture experiments suggest that xenogeneic cells may affect not only tumor cell growth and differentiation, but also, their interactions with immune system.

Future directions will include analysis of biomolecules secreted by chicken embryonic fibroblasts in paracrine fashion in co-culture. These biomolecules may have human homologs with potential utility in cancer treatment. We are also planning to explore this phenomenon further *in vivo* using animal models.

## Acknowledgments

We are indebted to Dr. Judith Luborsky for assistance in manuscript preparation. We would also like to acknowledge the generous contributions of Mr. and Mrs. Eugene and Shirley Deutsch, Mr. and Mrs. Mark and Maha Halabi Ditsch, Mr. George Ruwe, Clow Family Foundation, and Eisenbrandt Family Foundation.

## REFERENCES

[1] Robson M, Im S-A, Senkus E, Xu B, Domchek SM et al. Olaparib for metastatic breast cancer in patients with a germline brca mutation. New England Journal of Medicine 2017; 377: 523–533. DOI: 10.1056/NEJMoa1706450

[2] Audeh MW, Carmichael J, Penson RT, Friedlander M, Powell B et al. Oral poly(adp-ribose) polymerase inhibitor olaparib in patients with brca1 or brca2 mutations and recurrent ovarian cancer: A proof-of-concept trial. Lancet 2010; 376: 245–251. DOI: 10.1016/S0140-6736(10)60893-8

[3] Iqbal N, Iqbal N. Imatinib: A breakthrough of targeted therapy in cancer. Chemother Res Pract 2014; 2014: 357027. DOI: 10.1155/2014/357027

[4] Christodoulou I, Goulielmaki M, Devetzi M, Panagiotidis M, Koliakos G et al. Mesenchymal stem cells in preclinical cancer cytotherapy: A systematic review. Stem Cell Research & Therapy 2018; 9: 336. DOI: 10.1186/s13287-018-1078-8

[5] Liu T, Han C, Wang S, Fang P, Ma Z et al. Cancer-associated fibroblasts: An emerging target of anti-cancer immunotherapy. Journal of Hematology & Oncology 2019; 12: 86. DOI: 10.1186/s13045-019-0770-1

[6] Androulla NM, Lefkothea CP. Car t-cell therapy: A new era in cancer immunotherapy. Current Pharmaceutical Biotechnology 2018; 19: 5–18. DOI: http://dx.doi.org/10.2174/1389201019666180418095526

[7] Huang CP, Chen CC, Shyr CR. Xenogeneic cell therapy provides a novel potential therapeutic option for cancers by restoring tissue function, repairing cancer wound and reviving anti-tumor immune responses. Cancer Cell Int 2018; 18: 9. DOI: 10.1186/s12935-018-0501-7

[8] Strioga MM, Darinskas A, Pasukoniene V, Mlynska A, Ostapenko V et al. Xenogeneic therapeutic cancer vaccines as breakers of immune tolerance for clinical application: To use or not to use? Vaccine 2014; 32: 4015–4024. DOI: 10.1016/j.vaccine.2014.05.006

[9] Wei YQ, Wang QR, Zhao X, Yang L, Tian L et al. Immunotherapy of tumors with xenogeneic endothelial cells as a vaccine. Nat Med 2000; 6: 1160–1166. DOI: 10.1038/80506

[10] Tartour E, Mehtali M, Sastre-Garau X, Joyeux I, Mathiot C et al. Phase i clinical trial with il-2-transfected xenogeneic cells administered in subcutaneous metastatic tumours: Clinical and immunological findings. Br J Cancer 2000; 83: 1454–1461. DOI: 10.1054/bjoc.2000.1492

[11] Doi C, Maurya DK, Pyle MM, Troyer D, Tamura M. Cytotherapy with naive rat umbilical cord matrix stem cells significantly attenuates growth of murine pancreatic cancer cells and increases survival in syngeneic mice. Cytotherapy 2010; 12: 408–417. DOI: 10.3109/14653240903548194

[12] Huebener N, Fest S, Hilt K, Schramm A, Eggert A et al. Xenogeneic immunization with human tyrosine hydroxylase DNA vaccines suppresses growth of established neuroblastoma. Mol Cancer Ther 2009; 8: 2392–2401. DOI: 10.1158/1535-7163.MCT-09-0107

[13] Seledtsova GV, Shishkov AA, Kaschenko EA, Seledtsov VI. Xenogeneic cell-based vaccine therapy for colorectal cancer: Safety, association of clinical effects with vaccine-induced immune responses. Biomed Pharmacother 2016; 83: 1247–1252. DOI: 10.1016/j.biopha.2016.08.050

[14] Sohni A, Verfaillie CM. Mesenchymal stem cells migration homing and tracking. Stem Cells Int 2013; 2013: 130763. DOI: 10.1155/2013/130763

[15] Fiori ME, Di Franco S, Villanova L, Bianca P, Stassi G et al. Cancer-associated fibroblasts as abettors of tumor progression at the crossroads of emt and therapy resistance. Mol Cancer 2019; 18: 70. DOI: 10.1186/s12943-019-0994-2

[16] Liu T, Zhou L, Li D, Andl T, Zhang Y. Cancer-associated fibroblasts build and secure the tumor microenvironment. Front Cell Dev Biol 2019; 7: 60. DOI: 10.3389/fcell.2019.00060

[17] Lebleu VS, Kalluri R. A peek into cancer-associated fibroblasts: Origins, functions and translational impact. Dis Model Mech 2018; 11. DOI: 10.1242/dmm.029447

[18] Alkasalias T, Flaberg E, Kashuba V, Alexeyenko A, Pavlova T et al. Inhibition of tumor cell proliferation and motility by fibroblasts is both contact and soluble factor dependent. Proc Natl Acad Sci U S A 2014; 111: 17188–17193. DOI: 10.1073/pnas.1419554111

[19] Drexler HG, Uphoff CC. Mycoplasma contamination of cell cultures: Incidence, sources, effects, detection, elimination, prevention. Cytotechnology 2002; 39: 75–90. DOI: 10.1023/A:1022913015916

[20] Patro R, Duggal G, Love MI, Irizarry RA, Kingsford C. Salmon provides fast and bias-aware quantification of transcript expression. Nat Methods 2017; 14: 417–419. DOI: 10.1038/nmeth.4197

[21] Flaberg E, Markasz L, Petranyi G, Stuber G, Dicso F et al. High-throughput live-cell imaging reveals differential inhibition of tumor cell proliferation by human fibroblasts. International Journal of Cancer 2011; 128: 2793–2802. DOI: 10.1002/ijc.25612

[22] Hendrix MJC, Seftor EA, Seftor REB, Kasemeier-Kulesa J, Kulesa PM et al. Reprogramming metastatic tumour cells with embryonic microenvironments. Nature Reviews Cancer 2007; 7: 246. DOI: 10.1038/nrc2108

[23] Nixon NA, Blais N, Ernst S, Kollmannsberger C, Bebb G et al. Current landscape of immunotherapy in the treatment of solid tumours, with future opportunities and challenges. Curr Oncol 2018; 25: e373–e384. DOI: 10.3747/co.25.3840

[24] Bao Q, Zhao Y, Niess H, Conrad C, Schwarz B et al. Mesenchymal stem cell-based tumor-targeted gene therapy in gastrointestinal cancer. Stem Cells Dev 2012; 21: 2355–2363. DOI: 10.1089/scd.2012.0060

[25] Eyden B. Fibroblast phenotype plasticity: Relevance for understanding heterogeneity in “fibroblastic” tumors. Ultrastructural Pathology 2004; 28: 307–319. DOI: 10.1080/019131290882204

