## Supplementary Table 1 for "Xenogeneic Fibroblasts Inhibit The Growth Of Breast And Ovarian Cancer Cell Lines In Co-Culture"

Summary of data quality measures for Next Generation Sequencing results

| Sample | Raw Reads | Clean Reads | Effective Rate (%) | Error Rate (%) | Q20 (%) | Q30 (%) | GC Content (%) |
| --- | --- | --- | --- | --- | --- | --- | --- |
| Treated 1 | 42704286 | 42510515 | 99.55 | 0.03 | 95.25 | 88.81 | 48.58 |
| Treated 2 | 41128273 | 40920172 | 99.49 | 0.03 | 95.27 | 88.85 | 48.37 |
| Treated 3 | 40324878 | 40087851 | 99.41 | 0.02 | 95.32 | 88.94 | 48.50 |
| Control 1 | 40600290 | 40220009 | 99.06 | 0.03 | 95.32 | 88.86 | 48.33 |
| Control 2 | 41092608 | 40803301 | 99.3 | 0.02 | 95.67 | 89.53 | 48.45 |
| Control 3 | 45191393 | 44848640 | 99.24 | 0.03 | 94.75 | 87.84 | 48.33 |

Notes:

Clean bases: (Clean reads) * (sequence length), calculating in G. For paired-end sequencing like PE150, sequencing length equals 150, otherwise it equals 50 for sequencing like SE50.

Effective Rate (%): (Clean reads/Raw reads)*100%

Error rate: base error rate

Q20, Q30: (Base count of Phred value > 20 or 30) / (Total base count)

GC content: (G & C base count) / (Total base count)
